# Plant-insect chemical communication in ecological communities: an information theory perspective

**DOI:** 10.1101/2021.09.30.462509

**Authors:** Pengjuan Zu, Reinaldo García-García, Meredith Schuman, Serguei Saavedra, Carlos J. Melián

## Abstract

Cross-species communication, where signals are sent by one species and perceived by others, is one of the most intriguing types of communication that functionally links different species to form complex ecological networks. Yet, global changes and human activities can affect communication by increasing the fluctuations of species composition and phenology, altering signal profiles and intensity, and introducing noises. So far, most studies on cross-species communication have focused on a few specific species isolated from ecological communities. Scaling up investigations of cross-species communication to the community level is currently hampered by a lack of conceptual and practical methodologies. Here, we propose an interdisciplinary framework based on information theory to investigate mechanisms shaping cross-species communication at the community level. We use plants and insects, the cornerstones of most ecosystems, as a showcase; and focus on chemical communication as the key communication channel. We first introduce some basic concepts of information theory, then we illustrate information patterns in plant-insect chemical communication, followed by a further exploration of how to integrate information theory into ecological and evolutionary processes to form testable mechanistic hypotheses. We conclude by highlighting the importance of community-level information as a vehicle to better understand the maintenance of ecological systems, especially when facing rapid global changes.

## 1 Introduction

Communication is prevalent in nature. For example, honeybees waggle to send information that guide other bees in the colony (Von Frisch, 1974); birds sing to mate or alert others in the flock (Freeberg, 2008); flowers exhibit colors and scent that can attract pollinators or deter herbivores (Schoonhoven et al., 2005). Within these, the cross-kingdom (Plantae and Animalia) plant-insect communication is of extreme interest not only because of its ubiquity and fundamental roles in both natural and agricultural systems (Isbell et al., 2011; Ollerton, 2021; Potts et al., 2016; Seastedt and Crossley Jr, 1984; Strong et al., 1984), but also due to the extraordinarily diverse communication signals shaped by hundreds of millions of years of co-evolution (Ehrlich and Raven, 1964).

Chemical communication is one of the most ancient and pivotal means for plants and insects. In fact, chemistry underlies color (pigments), shape (genetic encoding in nucleic acids and chemical inducers such as hormones), and scent (volatile organic compounds, or VOCs). Yet, chemical signals often refer to the vast number of secondary metabolites produced by plants, with an estimate of 200,000 compounds that have been extracted and identified (Kessler and Kalske, 2018; Wink, 2010), that are not strictly required for plant growth and development, but play important defense and attraction functions (Ehrlich and Raven, 1964; Fraenkel, 1959; Pichersky and Gershenzon, 2002). Insects are very sensitive to many plant chemicals with their numerous chemical receptors (Hansson and Stensmyr, 2011; Kaupp, 2010; Schoonhoven et al., 2005) and depend on chemical signals for many fundamental activities such as foraging (Schiestl, 2010), mating (Alexander et al., 1997), and oviposition (Renwick and Chew, 1994). So far, most plant-insect chemical communication studies have focused on specific species, isolating them from the community context in which they are naturally embedded. However, scaling up plant-insect communication from species-level to community-level can be possible through the lens of communication systems (Shannon, 1948).

From an information theory perspective, communication is the process of information transferring from a sender who codes the information (e.g., visual, vocal, or olfactory) to a receiver who decodes the information (Box 1). Depending on how well the information is *reproduced* at the receiver’s end, we can quantify the clarity or ambiguity of the communication (Shannon, 1948). That is, the better a message is reproduced, the greater amount of information (clarity) it contains, and the lower its entropy (ambiguity, uncertainty). Box 1 summarizes key information quantities. Information theory was originally developed for man-made communication systems such as telecommunication, and later applied to a wide range of communication-related fields, for example, cybernetics (Gabor, 1954), cryptography (Ahlswede and Csiszár, 1993), linguistics (i Cancho and Solé, 2003; Zipf, 1949) and neurobiology (Sharpee et al., 2014). Information theory brings at least two important new features that make it a promising tool for various systems and scales. First, information theory translates diverse communication signals into information bits. Thus, it unites different signals by extracting their information content without being overwhelmed by their different identities. Second, information theory measures the reproducibility of information between two states based on probabilistic concepts (Boso and Tartakovsky, 2018). This nonparametric perspective allows the scaling of information across different dimensions.

### Box 1. Information transferring process in communications and formula of quantifying information in communication systems.

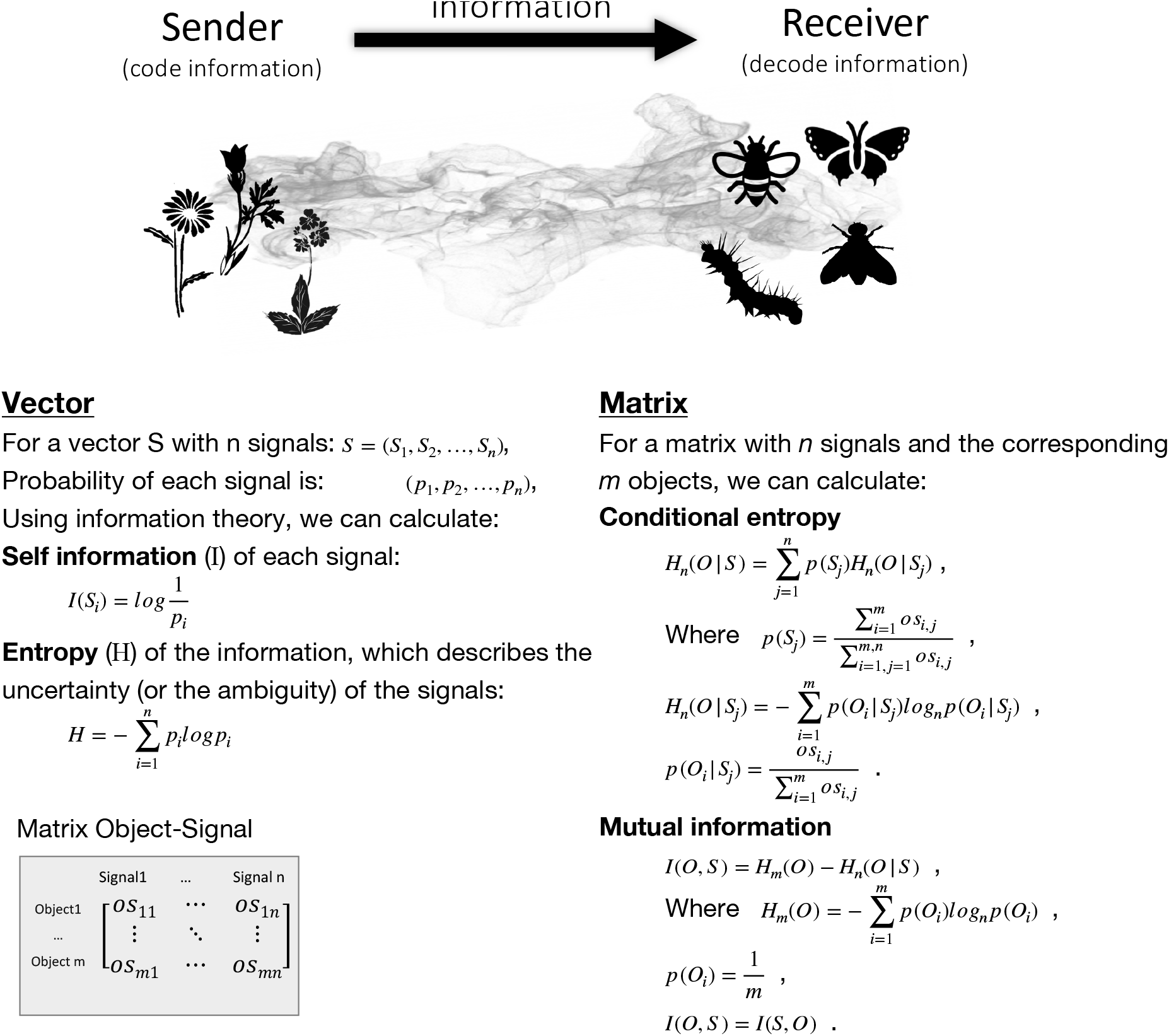

Not surprisingly, information theory has long been proposed as a conceptual and quantitative framework to study information transfer across ecological systems (Margalef, 1968*a*; O’Connor et al., 2019; Ulanowicz, 2001). However, concrete applications to plant-insect communication are just starting to emerge (Zu et al., 2020). In this paper, we illustrate the application of information theory as a vehicle to increase our understanding of plant-insect communication. In particular, we propose to use plant VOCs as communication signals, and explore the emerging information patterns of plant-insect chemical communication. We then discuss potential future directions to uncover the underlying mechanisms of plant-insect communication by the integration of information theory with ecological and evolutionary processes. Note that here we use “signals” in an information theory framework to simply refer to *all* plant VOCs that form the repository (regardless whether they have attractant or deterrent functions or not).

## 2 Patterns of plant-insect chemical communication

### 2.1 Zipf’s law

Zipf’s law (Zipf, 1932, 1949) in linguistics describes that the frequency of a word decays rapidly proportional to its rank, or frequency of usage, following a power law distribution (*p*(*r*) = *ar^−k^*, where *r* denotes rank). Taking the English language for example, “the” is the most frequent word in corpora (e.g., British National Corpus), occurring at a rate of 6% (according to WordCount: http://www.wordcount.org), followed by “of”, which only occurs half as often (3%). And there is a large number of words that occur in very low frequency, leading to a heavy tail distribution. The power-law decay distribution is also related to the Pareto principle (or 80-20 rule) known in social sciences (Pareto, 1964). Importantly, the power-law exponent is around *k* = 1 in many human languages (Yu et al., 2018) (e.g., English, German, Chinese), describing the speed of decay (i.e., the shape of this tailed distribution) and the redundancy of information in communication systems.

In terms of plants’ chemical language within ecological communities, plants emit different chemical “words” to their environment. Studies on plant VOCs have revealed that some VOCs are more frequent than others (Farré-Armengol et al., 2020; Knudsen et al., 2006), but the detailed information structures have not been explicitly studied. To explore the redundancy of plant chemical language and whether the frequency of its vocabulary (VOCs) follows Zipf’s law, we gathered the data from the only four community-level plant VOC studies so far, three of which focused on floral VOCs in plant-pollinator networks (Burkle and Runyon, 2019; Filella et al., 2013; Kantsa et al., 2017) and one on leaf VOCs in a plant-herbivore (caterpillar) network (Zu et al., 2020). Additionally, we also use a review study by Farré-Armengol et al. 2020 (Farré-Armengol et al., 2020), who compiled a floral VOC dataset from 305 species. In the review study, the authors also categorized chemical groups of VOCs, families of the plant species, and pollination systems of these plant species.

By analysing the distributions of VOCs in these studies and in different categories (chemical groups, plant families, pollination groups), we found that most cases follow a heavy-tail distribution where a few VOCs are predominant whereas many other VOCs occur rarely (Fig. 1). Only in 5 cases (out of the 16 examined cases and categories), a power-law distribution fits either the whole data (in one community study *PV_k*, and the group of N-, S-containing compounds, Fig. 1, Table 1) or part of the data (in Orchidaceae, insect-pollinated flowers, and the overall review data, Fig. 1, Table 1). The fitting was performed using the R package “poweRlaw” (Clauset et al., 2009; Gillespie, 2017). Many of the left cases are power-law like but cannot be statistically supported. Indeed, power law distribution is thought to be common but not easy to achieve a strict fit to many empirical data (Clauset et al., 2009) due to limited sample size and sample bias on abundant vs. rare signals.

**Fig. 1:**
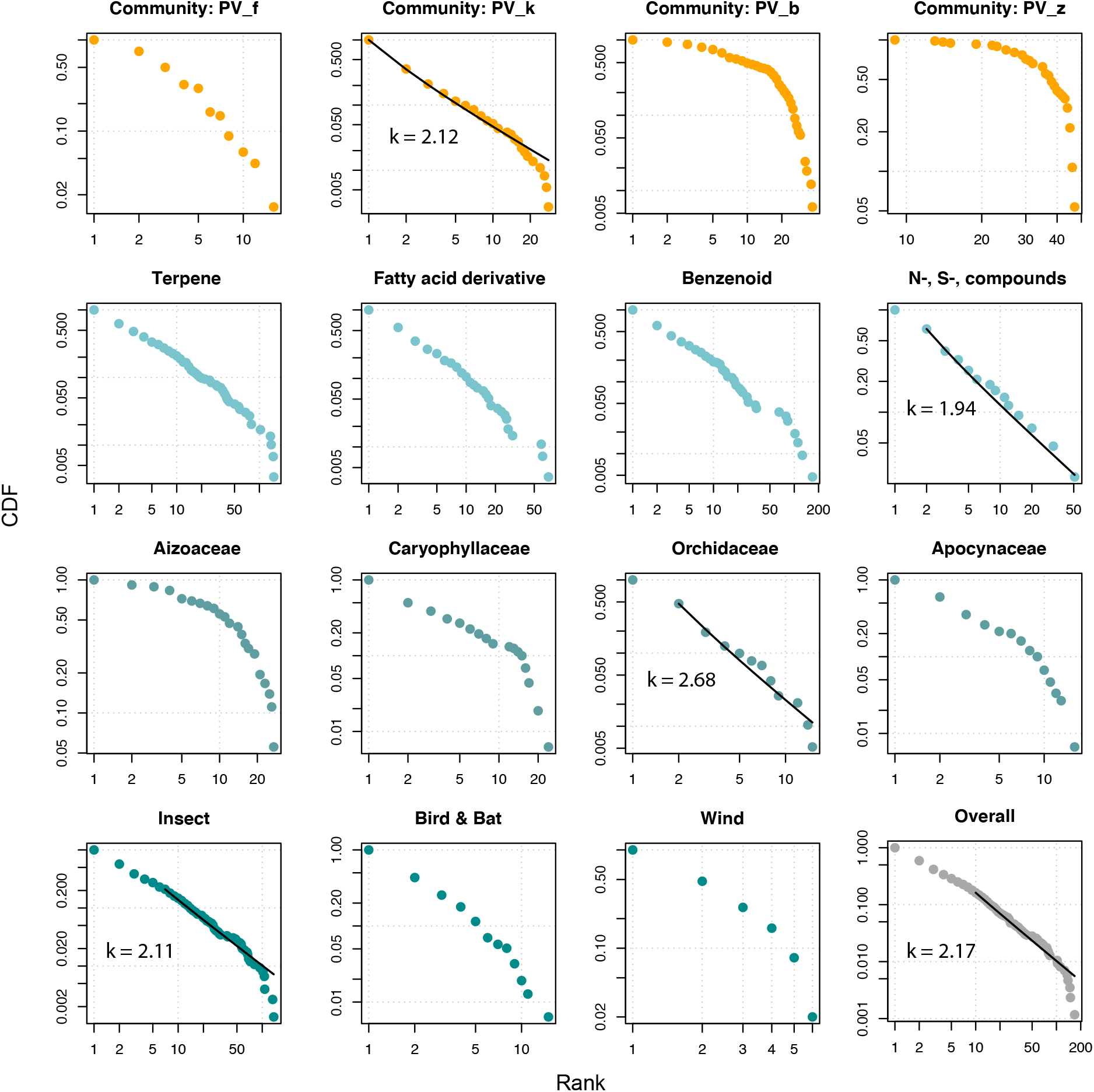
VOC frequency vs rank (log-log scale) from four community level study papers (top row) and a review paper (2nd to 4th rows) that summarised VOC frequency in different chemical groups (2nd row), four representatives of plant families (3rd row), and plants with different pollination groups and the overall review data (4th row). Black lines in the cases of “Community: PV_k”, “N-, S-, compounds”, “Orchidaceae”, “Overall” indicate that data can be described by power low distribution (k values represent the exponent of the power law distribution *p*(*r*) = *ar^−k^*), although log normal distribution can describe the data similarly well (tests see Supplementary Table S1). N-, S-, compounds: nitrogen or sulphur containing volatile compounds. Number of sampled plant species (N_plant) and number of VOCs (N_VOC) in each case can be found in Table 1.

**Table 1.**
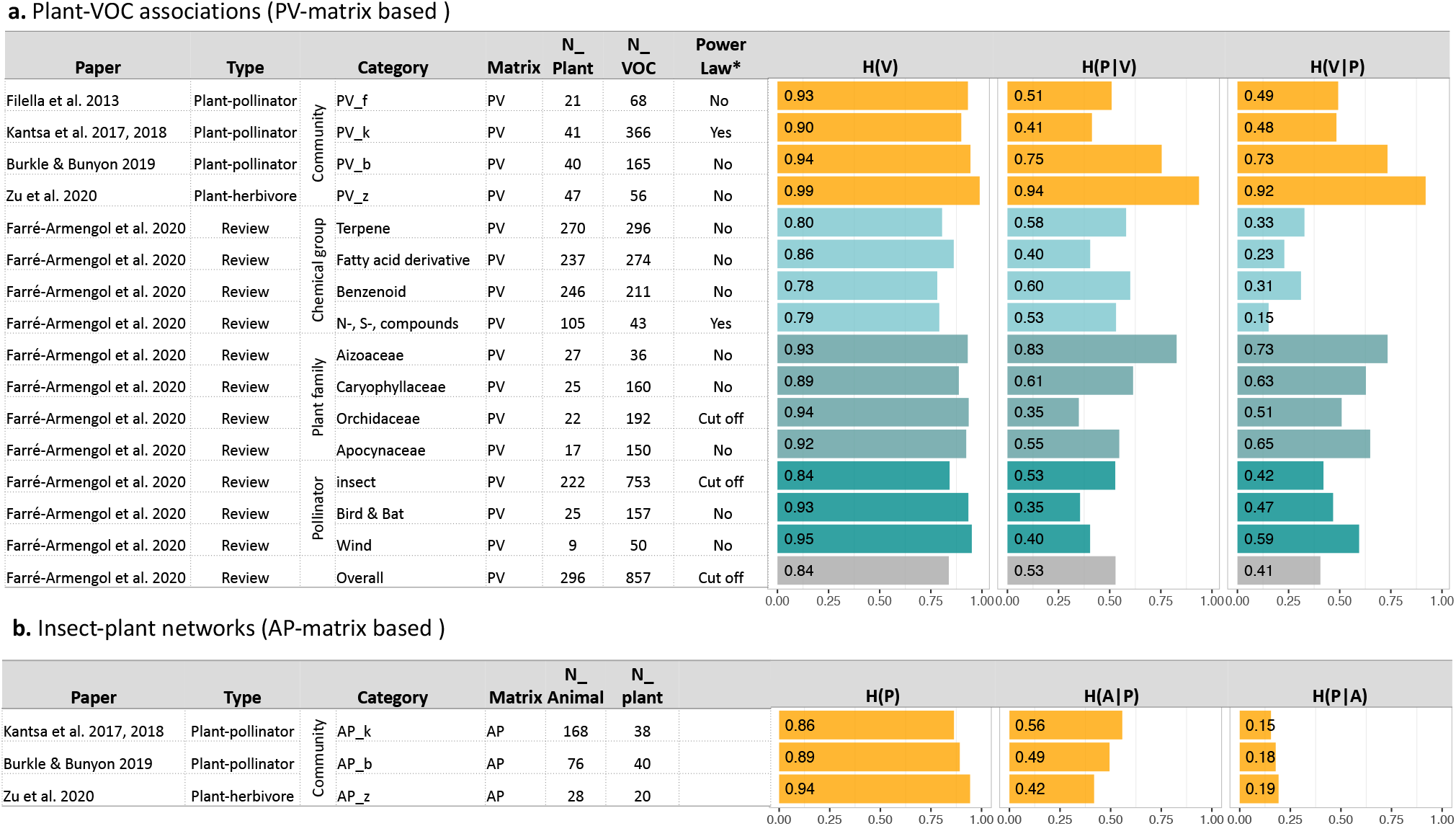
Entropy measures based on **a)** Plant-VOC associations (PV-matrix based) and **b)** Insect-plant networks (AP-matrix based) in the four community level studies papers and a review paper that compiled plant-VOC publications. H(V) entropy of VOCs. *H*(*|*) conditional entropy. Calculation formulas see Box 1.

In these 5 cases, the slopes of the distribution range from *k* = 1.94 to *k* = 2.68, which is steeper than observed in human languages. These patterns suggest that plant VOC languages may be more redundant than human language. Following information theory, the entropy of VOCs in all these cases is very high, ranging from 0.78 to 0.99 (*H*(*V*) in Table 1a). Note that throughout the whole paper, we used standardised entropy so that it ranges between 0 and 1, where the higher the value, the higher the uncertainty. Entropy calculation formulas can be found in Box 1.

### 2.2 Coding, decoding and interactions

In the previous section, we only focus on the putative signals (VOCs) themselves. In this section, we place these into the context of communication: the coding and decoding of signals, and species interactions. Following the work by Zu et al. (2020), we can use how VOCs (V) are associated with plants (P) in a community to describe plant coding process (PV-matrix), how VOCs are associated with insects (or animals, A) to describe insect decoding process (AV-matrix), and associations of animals with plants (AP-matrix) for insect-plant interactions. Empirically, to screen how each insect reacts to each of the VOCs in the community requires extensive experimental manipulations under both controlled an natural conditions. Therefore, it is challenging to generate the whole insect decoding matrix from field and experimental work.

From the four community-level plant VOC studies which documented the PV-matrix, three of them (Burkle and Runyon, 2019; Kantsa et al., 2019; Zu et al., 2020) also document insect-plant interaction networks (AP-matrix) in the same community. We found that plant coding patterns vary from community to community (Fig. 2, Table 1). In the case of the leaf VOC community (PV_z), VOCs are more intensively connected or shared among plants (*H*(*P*|*V*) = 0.94), whereas in two cases of flower VOCs (PV_f, PV_k), there are more unique VOCs (*H*(*P*|*V*) = 0.41, 0.51 respectively), and the third case of flower VOCs is in between (*H*(*P*|*V*) = 0.75). We also analyzed plant coding patterns based on the review study, and found these to vary from category to category (Table 1a), with the overall (from the whole dataset) value of around 0.5 indicating some specialization (*H*(*P*|*V*) = 0.53, *H*(*V*|*P*) = 0.41, Table 1a).

**Fig. 2:**
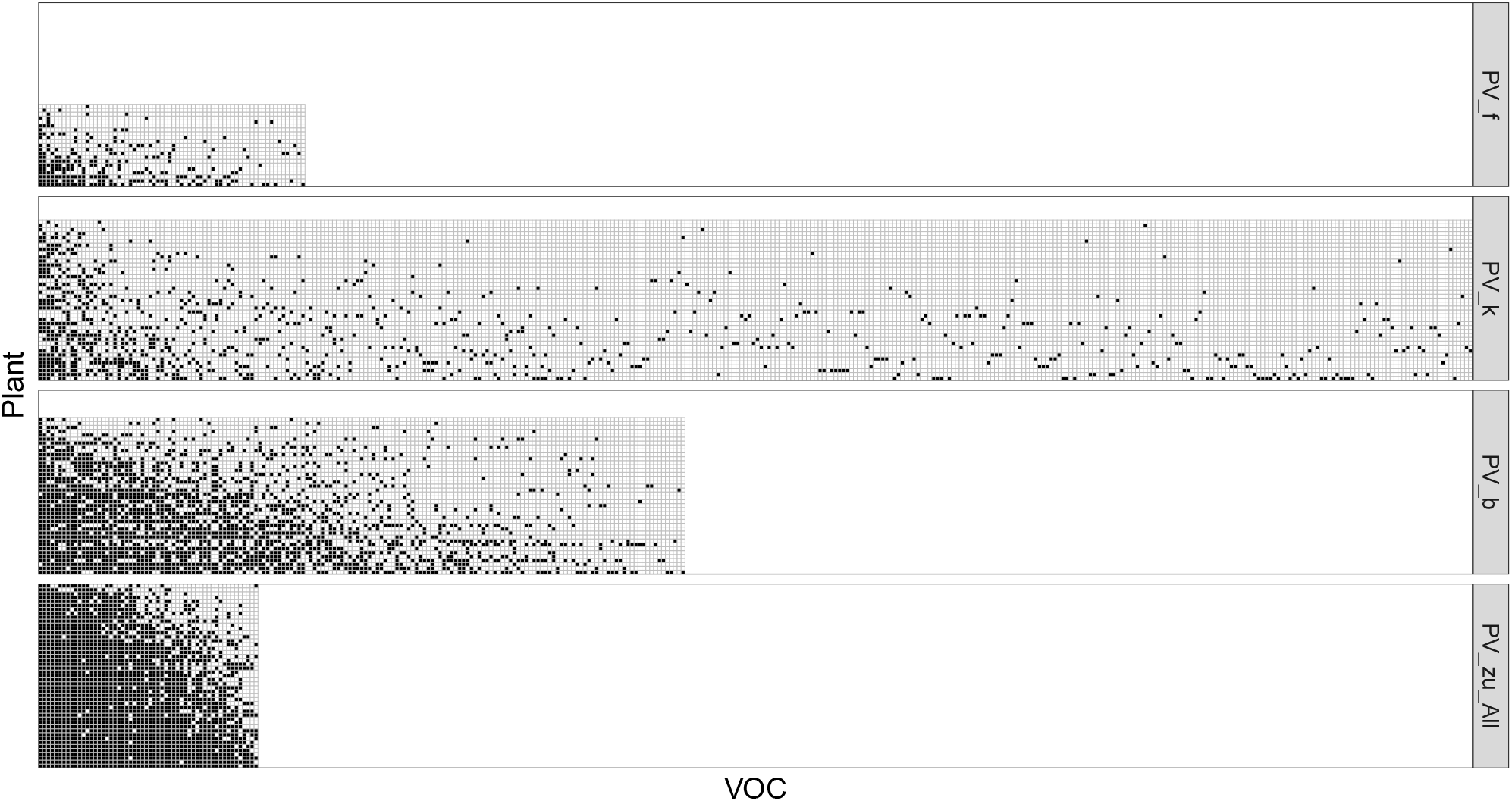
Patterns of plant-VOC associations (PV-matrix) in the four community studies (details see top four rows in Table 1a). Each row and column represents plant species and VOCs respectively. Black and white squares show that a given VOC was present or absent in a given plant, respectively.

**Fig. 3:**
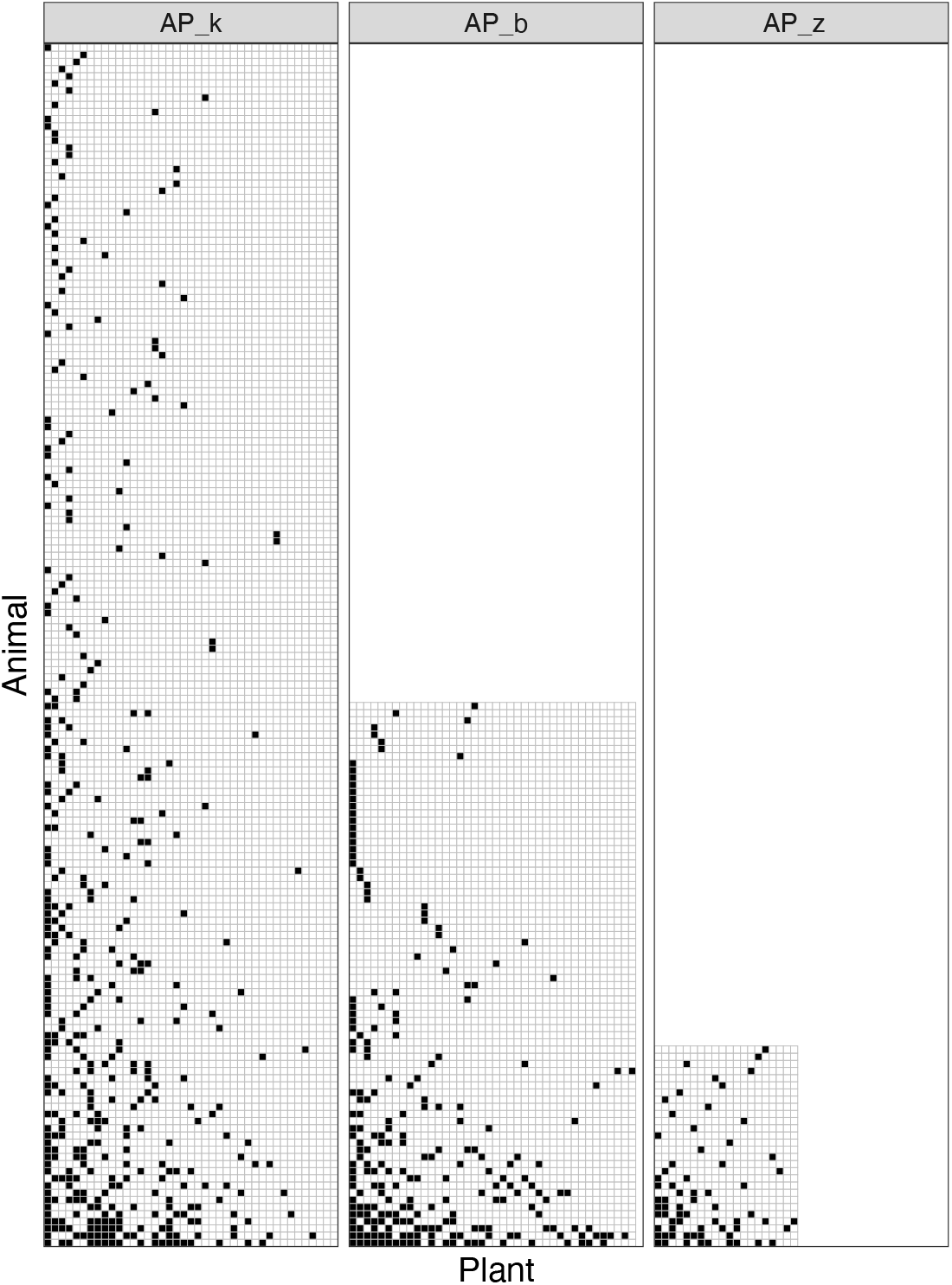
Patterns of insect-plant interactions (AP-matrix) in the three community studies (details see Table 1b). Each row and column represents insect species and plant species respectively. Black and white squares show that a given insect was, or was not observed feeding on a given plant, respectively

The three insect-plant interactions exhibit specialised patterns for both mutualistic pollinator-plant interactions and antagonistic herbivore-plant interactions as revealed by entropy values much less than 1 (*H*(*A*|*P*) around 0.5, *H*(*P*|*A*) around 0.2, Table 1b). The asymmetric values of *H*(*A*|*P*) and *H*(*P*|*A*) indicate that insects are more specialized on plants than plants specialized on insects (*H*(*P*|*A*) are lower).

## 3 From patterns to mechanisms

We have shown the information structures of plant-insect chemical communications as revealed by the apparent power-law behavior in some cases (but where these conclusions are limited by limited data availability), and by the ubiquitous decrease in entropy of VOC associations to plants as opposed to total entropy of VOCs. The latter is not surprising, as not all plants produce all VOCs. Interestingly, the conditional entropy of VOCs (*H*(*P*|*V*)) involved in plant-herbivore interactions seems to be generally higher than the entropy of those involved in plant-pollinator interactions; but this is not reflected in the conditional entropy of plant-animal associations (*H*(*A*|*P*)) - which is similar among these interaction types. This suggests that plants try to communicate more clearly with their pollinators than with their herbivores, while herbivores must try harder to decode information from their host plants than pollinators.

Why are plant-insect chemical communications structured in the way they are and not otherwise? In this section, we highlight two directions of theoretical frameworks to explore the potential mechanisms driving these chemical communication patterns between plants and insects. Recall the difficulties of building a complete decoding matrix (how each insect decodes each of the VOCs) empirically. Here we theoretically construct insect decoding from two approaches: 1) using a top-down logic assuming that the insect can decode all the VOCs from a plant as long as they can feed on that plant, that is, AV = AP * PV; and 2) use a bottom-up logic by hypothesizing the relationships between VOC abundance and functionality (i.e., effects on insects). In both directions, we integrate information theory into ecological and evolutionary theories to generate testable hypotheses.

### 3.1 Top-down: Interactions shape communication

This direction follows the inspiration of studies exploring how Zipf’s law patterns emerge in linguistics. Various hypotheses have been proposed to explain this intriguing pattern in human languages. Among these, the “least effort” hypothesis (i Cancho and Solé, 2003; Zipf, 1949) is a compelling hypothesis that nicely recovers the Zipf’s law distribution as a natural consequence from the conflicting interests between speakers (who are thought to aim for “brevity and phonological reduction” to minimize their effort to speak) and listeners (who should desire “explicitness and clarity” to minimize their effort to understand). The successful application of information theory in studying the structure and emergence of human language has inspired studies on animal vocal communication (e.g., reviews in Kershenbaum et al. (2021); McCowan et al. (2008); McCOWAN et al. (1999)), and indicates its potential to be extended in studying the chemical “language” in plant-insect communities (Zu et al., 2020).

Indeed, Zu et al. (2020) aimed to borrow the framework to test whether plant-herbivore chemical communication patterns can be explained by conflicting interests of plants (speakers) and herbivores (listeners). It is well-known that the chemical arms race between plants and herbivores has been playing out for hundreds of millions of years, where herbivores must constantly adapt to plant defense chemicals and plants need to keep producing novel chemicals (Ehrlich and Raven, 1964). Zu and colleagues (Zu et al., 2020) translated this chemical arms race into an information arms race: plants code the (chemical) information in a way (changes in PV-matrix) to make the decoding (AV given by AP * PV) difficult (high entropy) for herbivores, whereas herbivores interact with plants in a way (changes in AP-matrix) to make the decoding easier (low entropy). With repeated cycles of information optimisation by plants and herbivores, they found that an equilibrium stage is reached in which a plant-VOC redundancy matrix and herbivore-plant specialization matrix have emerged. These information patterns at the equilibrium stage matched their field data collected in a tropical dry forest.

This is a successful case of integrating other known evolutionary and ecological theories into information theory to build a framework that aims to disentangle mechanisms driving plant-insect interactions and communications at the community level. In general, there are three steps to construct communication frameworks for a given system. First, we must define who are the speakers and who are the listeners. Second, we must define proper fitness functions for both parties based on their ecological and evolutionary relationships. Third, we must define the rules of optimization for each party in the information game. Both the second and the third steps require very careful examination by integrating evolutionary and ecological processes. We also want to emphasize that the “fitness” function at the community level is not the same as the “fitness” biologists normally used for measuring survival and reproductive success, but rather refers to broader benefits which can be assessed at the level of communities rather than individuals. This corresponds as well to the broader use of the term “signal” introduced at the beginning of this paper. This approach is compatible with the evaluation of individual-level fitness, but aims at criteria which can be used at the community level. We hope more studies will be inspired in this information perspective to bring us new insights in more plant-insect chemical communication types (e.g., plant-pollinator, plant-herbivore-parasitoid).

### 3.2 Bottom-up: Connect VOC frequency with “information functionality”

Besides the top-down method, in this section, we want to briefly point out another potential angel: bottom-up, connecting VOC frequency with information functionality to study plant-insect communication. Given there are rare and common VOCs, it is natural to ask whether rare VOCs or common VOCs can be decoded more easily by insects (functionality) and how much information can be gained by decoding the VOCs (information).

Shannon’s original mathematical theory of communication (Shannon, 1948) was mostly based on the “Inverse Relationship Principle” (D’Alfonso, 2011) which states that the less probable a signal is, the more information it bears. In our plant-insect communication systems, insects face an environment with VOCs in different frequencies. On the one hand, specializing in decoding rare VOCs brings advantages if substantial information is thereby gained to access a niche that can only be accessed by specialists. However, specialization comes with risks because rare VOCs (emitted by only one plant species or a few phenotypes) can go extinct more easily. On the other hand, decoding common VOCs is likely to be safer but less informative, meaning that insects might need to decode more signals to gather enough information to identify their plant hosts. Which strategy do insects use and how will different strategies affect the interaction network structure? For example, will positive-frequency dependent decoding structure, therefore decoding preferentially common VOCs, lead to more generalized plant-insect interactions; whereas negative-frequency dependent, decoding preferentially rare VOCs, results in more specialized interactions?

In this framework, one can hypothesize different relationships between VOC frequency and its information functionality (VA-matrix), to test how they give rise to plant-insect interaction network structures (using *PA* = *PV* * *VA*). Specifically, information functionality can be modelled with two elements, one is how easy a VOC can be decoded (*V A_func*), the other is how much information that can be gained by decoding the VOC (*V_info*). Therefore, the information functionality matrix (VA) can be treated as an information-weighted decoding matrix, generated by (*VA* = *V_info* * *V A_func*). We can use a drift process mimicking frequency-independent decoding which might be used as a test for the strength of frequency-dependent decoding occurring in the empirical data.

## Conclusion

We have found from the limited studies that plant VOCs frequencies follow a heavy-tail distribution with a few predominant compounds and many rare compounds; all the three documented plant-insect interactions are highly specialized, whereas plant-VOC coding patterns seem to vary from case to case with some plant communities coding the signals more redundantly than others. We also provide a top-down and a bottom-up direction for constructing theoretical frameworks to integrate information theory in studying plant-insect chemical communication. by bridging top vs. bottom up frameworks, we hope we can draw attention and inspire more studies on the information perspective of trophic networks in ecosystems.

In the face of more serious and more frequent threats to ecological systems, plant-insect interactions are vulnerable, yet particularly important aspects of ecosystem functioning: insect pollinators are vital for agricultural productivity as well as for the survival of wild plant communities (Ollerton, 2021; Wei et al., 2021); herbivores support all higher trophic levels which depend on plants (Harvey et al., 2003; Moreira et al., 2016; Price et al., 1980), and their predators and parasitoids perform pest control services (Schmidt et al., 2003). We still struggle to control these interactions with relatively crude tools which carry heavy collateral damage, such as physically uprooting and transporting communities of pollinators to agricultural fields, and broad prescribed spraying of pesticides. Integrating information theory has the potential to help us gain a better understanding of the underlying mechanisms, and thus promises novel insights into challenging situations both from fundamental to applied ones. For example, novel chemicals of exotic plants have been suggested to be a main mechanism for invading the local communities (Cappuccino and Arnason, 2006; Macel et al., 2014). If we put it in the information perspective, we can ask what are the effects of invasive species on the information landscape of the current community; whether there are some information structures more resilient than others. Moreover, how will climate change, pollution (including pesticide) and habitat fragmentation affect communication signal profiles, dynamics, and signal-to-noise ratios and thus rewire plant-insect networks? Overall, understanding the effects of communication on species interactions and vice versa has the potential to improve our assessment of the impact of anthropogenic disturbances on ecosystems, and assist us for conservation and restoration practices. For instance, instead of focusing on taxonomy diversity, chemical diversity and chemical information structure may be shown to be more crucial and informative in affecting ecosystem biodiversity (Schuman et al., 2016).

Information approaches can provide the scalability to integrate further dimensions to achieve a better and potentially more unified understanding of information flows and their effects in ecosystems. For instance, we can expand these frameworks by including different sets of volatiles (e.g., from flowers, leaves and roots); different sets of partners and across trophic levels (e.g., pollinators, microbes, parasitoids and predators); and even different information forms (e.g., visual, vocal). Information theory abstracts all kinds of signals (or functional traits) as information, and thus has the potential to gather the multi-layer functional maps into a unified framework. In summary, integrating information theory to existing theories in ecology and evolution has the potential to unveil central biological mechanisms driving the formation and maintenance of the functionality of species interactions and thus of entire ecosystems (Kessler and Kalske, 2018; Margalef, 1968*b*; O’Connor et al., 2019).

## Acknowledgments

PZ acknowledges the Swiss National Science Foundation (SNSF) Spark grant and SNSF Prima grant, under number CRSK-3_196506 and PR00P3_193237 respectively. MCS acknowledges the support of the University of Zurich University Research Priority Program on Global Change and Biodiversity and membership in the US NSF-funded Biology Integration Institute BII-Implementation: The causes and consequences of plant biodiversity across scales in a rapidly changing world (Award Number:2021898). Funding to SS was provided by NSF grant No. DEB-2024349.

## Supplementary Materials

**Table S1**. Statistical results of power law tests for the 16 cases in Fig. 1.

## Author contributions

PZ initiated and performed the study. All authors contributed with ideas and wrote the manuscript.

## Data accessibility

The data and R code supporting the results can be found at https://github.com/PengjuanZu/Zu_etal_Plant-insect_InformationTheory.

## Bibliography

Ahlswede, R. and Csiszár, I. 1993. Common randomness in information theory and cryptography. i. secret sharing. IEEE Transactions on Information Theory 39:1121–1132.

Alexander, R. D., Marshall, D. C., and Cooley, J. R. 1997. Evolutionary perspectives on insect mating. The evolution of mating systems in insects and arachnids Pages 4–31.

Boso, F. and Tartakovsky, D. M. 2018. Information-theoretic approach to bidirectional scaling. Water Resources Research 54:4916–4928.

Burkle, L. A. and Runyon, J. B. 2019. Floral volatiles structure plant–pollinator interactions in a diverse community across the growing season. Functional Ecology 33:2116–2129.

Cappuccino, N. and Arnason, J. T. 2006. Novel chemistry of invasive exotic plants. Biology letters 2:189–193.

Clauset, A., Shalizi, C. R., and Newman, M. E. 2009. Power-law distributions in empirical data. SIAM review 51:661–703.

D’Alfonso, S. 2011. On quantifying semantic information. Information 2:61–101.

Ehrlich, P. R. and Raven, P. H. 1964. Butterflies and plants: a study in coevolution. Evolution 18:586–608.

Farré-Armengol, G., Fernández-Martínez, M., Filella, I., Junker, R. R., and Peñuelas, J. 2020. Deciphering the biotic and climatic factors that influence floral scents: a systematic review of floral volatile emissions. Frontiers in plant science 11:1154.

Filella, I., Primante, C., Llusia, J., González, A. M. M., Seco, R., Farré-Armengol, G., Rodrigo, A., Bosch, J., and Penuelas, J. 2013. Floral advertisement scent in a changing plant-pollinators market. Scientific Reports 3:3434.

Fraenkel, G. S. 1959. The raison d’etre of secondary plant substances. Science Pages 1466–1470.

Freeberg, T. M. 2008. Complexity in the chick-a-dee call of carolina chickadees (poecile carolinensis): associations of context and signaler behavior to call structure. The Auk 125:896–907.

Gabor, D. 1954. Communication theory and cybernetics. Transactions of the IRE Professional Group on Circuit Theory Pages 19–31.

Gillespie, C. S. 2017. Estimating the number of casualties in the american indian war: a bayesian analysis using the power law distribution. The Annals of Applied Statistics 11:2357–2374.

Hansson, B. S. and Stensmyr, M. C. 2011. Evolution of insect olfaction. Neuron 72:698–711.

Harvey, J. A., Van Dam, N. M., and Gols, R. 2003. Interactions over four trophic levels: foodplant quality affects development of a hyperparasitoid as mediated through a herbivore and its primary parasitoid. Journal of Animal Ecology 72:520–531.

i Cancho, R. F. and Solé, R. V. 2003. Least effort and the origins of scaling in human language. Proceedings of the National Academy of Sciences 100:788–791.

Isbell, F., Calcagno, V., Hector, A., Connolly, J., Harpole, W. S., Reich, P. B., Scherer-Lorenzen, M., Schmid, B., Tilman, D., Van Ruijven, J., et al. 2011. High plant diversity is needed to maintain ecosystem services. Nature 477:199–202.

Kantsa, A., Raguso, R. A., Dyer, A. G., Sgardelis, S. P., Olesen, J. M., and Petanidou, T. 2017. Community-wide integration of floral colour and scent in a mediterranean scrubland. Nature Ecology & Evolution 1:1502.

Kantsa, A., Raguso, R. A., Lekkas, T., Kalantzi, O.-I., and Petanidou, T. 2019. Floral volatiles and visitors: A meta-network of associations in a natural community. Journal of Ecology.

Kaupp, U. B. 2010. Olfactory signalling in vertebrates and insects: differences and commonalities. Nature Reviews Neuroscience 11:188.

Kershenbaum, A., Demartsev, V., Gammon, D. E., Geffen, E., Gustison, M. L., Ilany, A., and Lameira, A. R. 2021. Shannon entropy as a robust estimator of zipf’s law in animal vocal communication repertoires. Methods in Ecology and Evolution 12:553–564.

Kessler, A. and Kalske, A. 2018. Plant secondary metabolite diversity and species interactions. Annual Review of Ecology, Evolution, and Systematics 49:115–138.

Knudsen, J. T., Eriksson, R., Gershenzon, J., and Ståhl, B. 2006. Diversity and distribution of floral scent. The Botanical Review 72:1.

Macel, M., de Vos, R. C., Jansen, J. J., van der Putten, W. H., and van Dam, N. M. 2014. Novel chemistry of invasive plants: exotic species have more unique metabolomic profiles than native congeners. Ecology and Evolution 4:2777–2786.

Margalef, R. 1968*a*. Perspectives in ecological theory.

Margalef, R., 1968*b*. Perspectives in Ecological Theory. University of Chicago Press, Chicago.

McCowan, B., Doyle, L., Hanser, S., Kaufman, A., and Burgess, C., 2008. Detection and estimation of complexity and contextual flexibility in nonhuman animal communication systems. MIT Press: Cambridge, MA, USA.

McCOWAN, B., Hanser, S. F., and Doyle, L. R. 1999. Quantitative tools for comparing animal communication systems: information theory applied to bottlenose dolphin whistle repertoires. Animal behaviour 57:409–419.

Moreira, X., Abdala-Roberts, L., Rasmann, S., Castagneyrol, B., and Mooney, K. A. 2016. Plant diversity effects on insect herbivores and their natural enemies: current thinking, recent findings, and future directions. Current opinion in insect science 14:1–7.

O’Connor, M. I., Pennell, M., Altermatt, F., Matthews, B., Melian, C., and Gonzalez, A. 2019. Principles of ecology revisited: integrating information and ecological theories for a more unified science. Frontiers in Ecology and Evolution 7:219.

Ollerton, J., 2021. Pollinators and Pollination: Nature and Society. Pelagic Publishing Ltd.

Pareto, V., 1964. Cours d’économie politique, volume 1. Librairie Droz.

Pichersky, E. and Gershenzon, J. 2002. The formation and function of plant volatiles: perfumes for pollinator attraction and defense. Current opinion in plant biology 5:237–243.

Potts, S. G., Imperatriz-Fonseca, V., Ngo, H. T., Aizen, M. A., Biesmeijer, J. C., Breeze, T. D., Dicks, L. V., Garibaldi, L. A., Hill, R., Settele, J., et al. 2016. Safeguarding pollinators and their values to human well-being. Nature 540:220–229.

Price, P. W., Bouton, C. E., Gross, P., McPheron, B. A., Thompson, J. N., and Weis, A. E. 1980. Interactions among three trophic levels: influence of plants on interactions between insect herbivores and natural enemies. Annual review of Ecology and Systematics 11:41–65.

Renwick, J. and Chew, F. 1994. Oviposition behavior in lepidoptera. Annual Review of Entomology 39:377–400.

Schiestl, F. P. 2010. The evolution of floral scent and insect chemical communication. Ecology Letters 13:643–656.

Schmidt, M. H., Lauer, A., Purtauf, T., Thies, C., Schaefer, M., and Tscharntke, T. 2003. Relative importance of predators and parasitoids for cereal aphid control. Proceedings of the Royal Society of London. Series B: Biological Sciences 270:1905–1909.

Schoonhoven, L. M., Van Loon, B., van Loon, J. J., and Dicke, M., 2005. Insect-plant biology. Oxford University Press on Demand.

Schuman, M. C., van Dam, N. M., Beran, F., and Harpole, W. S. 2016. How does plant chemical diversity contribute to biodiversity at higher trophic levels? Current opinion in insect science 14:46–55.

Seastedt, T. and Crossley Jr, D. 1984. The influence of arthropods on ecosystems. Bioscience 34:157–161.

Shannon, C. E. 1948. A mathematical theory of communication. Bell System Technical Journal 27:379–423.

Sharpee, T. O., Calhoun, A. J., and Chalasani, S. H. 2014. Information theory of adaptation in neurons, behavior, and mood. Current opinion in neurobiology 25:47–53.

Strong, D. R., Lawton, J. H., Southwood, S. R., et al., 1984. Insects on plants. Community patterns and mechanisms. Blackwell Scientific Publicatons.

Ulanowicz, R. E. 2001. Information theory in ecology. Computers & Chemistry 25:393–399.

Von Frisch, K. 1974. Decoding the language of the bee. Science 185:663–668.

Wei, N., Kaczorowski, R. L., Arceo-Gómez, G., O’Neill, E. M., Hayes, R. A., and Ashman, T.-L. 2021. Pollinators contribute to the maintenance of flowering plant diversity. Nature Pages 1–5.

Wink, M., 2010. Annual plant reviews, functions and biotechnology of plant secondary metabolites, volume 39. John Wiley & Sons.

Yu, S., Xu, C., and Liu, H. 2018. Zipf’s law in 50 languages: its structural pattern, linguistic interpretation, and cognitive motivation. arXiv preprint arXiv:1807.01855.

Zipf, G. 1932. Selected studies of the principle of relative frequency in language..

Zipf, G. K. 1949. Human behavior and the principle of least effort: an introd. to human ecology.

Zu, P., Boege, K., Del-Val, E., Schuman, M. C., Stevenson, P. C., Zaldivar-Riverón, A., and Saavedra, S. 2020. Information arms race explains plant-herbivore chemical communication in ecological communities. Science 368:1377–1381.

